# DRT111/SFPS splicing factor controls ABA sensitivity in Arabidopsis seed development and germination

**DOI:** 10.1101/2020.02.07.939421

**Authors:** Paola Punzo, Alessandra Ruggiero, Marco Possenti, Giorgio Perrella, Roberta Nurcato, Antonello Costa, Giorgio Morelli, Stefania Grillo, Giorgia Batelli

**Affiliations:** National Research Council of Italy, Institute of Biosciences and Bioresources (CNR-IBBR), Via Università 133, Portici (NA), Italy; Research Centre for Genomics and Bioinformatics, Council for Agricultural Research and Economics (CREA-GB), Via Ardeatina 546, 00178 Rome, Italy; ENEA Italian National Agency for New Technologies, Energy and Sustainable Economic Development, Trisaia Research Center, S.S. 106 Ionica, km 419+500 - 75026 Rotondella (MT), Italy

**Keywords:** *Arabidopsis thaliana*, pre-mRNA splicing, abscisic acid, light signaling, seed germination, transcriptomics

## Abstract

RNA splicing is a fundamental mechanism contributing to the definition of the cellular protein population in any given environmental condition. DRT111/SFPS is a splicing factor previously shown to interact with phytochromeB and characterized for its role in splicing of pre-mRNAs involved in photomorphogenesis. Here, we show that DRT111 interacts with Arabidopsis Splicing Factor 1 (SF1), involved in 3’ splicing site recognition. Double and triple mutant analysis shows that DRT111 controls splicing of *ABI3* and acts upstream of the splicing factor SUPPRESSOR OF ABI3-5 (SUA). *DRT111* is highly expressed in seeds and stomata of *Arabidopsis* and is induced by long-term treatments with polyethylene glycol and ABA. *DRT111* knock-out mutants are defective in ABA-induced stomatal closure and are hypersensitive to ABA during seed germination. Conversely, *DRT111* over-expressing plants show ABA hyposensitive seed germination. RNAseq experiments show that in dry seeds, *DRT111* controls expression and splicing of genes involved in response to osmotic stress and ABA, light signaling and mRNA splicing, including targets of ABSCISIC ACID INSENSITIVE3 (ABI3) and PHYTOCHROME INTERACTING FACTORs (PIFs). Consistently, expression of the germination inhibitor *SOMNUS,* induced by ABI3 and PIF1 is up-regulated in imbibed seeds of *drt111-2* mutants. Altogether, these results indicate that *DRT111* controls sensitivity to abscisic acid (ABA) during seed development, germination and stomatal movements and constitutes a point of integration of the ABA- and light-regulated pathways to control seed germination.

**One Sentence Summary:** Arabidopsis splicing factor *DRT111/SFPS* is required for ABA-mediated responses in seeds

## Introduction

The phytohormone abscisic acid (ABA) regulates physiological and developmental processes, including stress responses, seed development and germination.

Perhaps the most well defined mechanism mediated by ABA is induction of stomatal closure.

In plants subjected to hyperosmotic stress, ABA is synthesized predominantly in leaf vascular tissues and guard cells. Here, ABA activates a signalling pathway that coordinately modulates activity of membrane located transporters, leading to efflux of solutes. The consequent reduction of turgor of guard cells causes stomatal closure, thus reducing evapotranspiration in abiotic stress conditions (Bauer et al.,2013; Kuromori et al.., 2018; Nambara and Marion-Poll, 2005; Qin and Zeevaart, 1999; Schroeder et al., 2001).

In seeds, ABA induces maturation, dormancy and plays a key role during germination. Transcription factors such as LEAFY COTYLEDON1 and 2 (LEC1 and LEC2), FUSCA3 (FUS3) and ABSCISIC ACID INSENSITIVE3 (ABI3) are involved in reserve accumulation and inhibition of premature germination (Santos-Mendoza et al., 2008, Monke et al., 2012, Yan and Zhen, 2017). At early stages of seed maturation, *LEC1/2* and *FUS3* are expressed to prevent germination of the developing embryo, whereas *ABI3* expression is maintained at high levels until final maturation stages (Perruc et al., 2007). In this phase, ABI3 and LEC1 regulate expression of genes involved in storage reserve accumulation and acquisition of desiccation tolerance, such as late embryogenesis abundant (LEA) proteins (Parcy et al., 1994).

In addition, ABA prevents germination by inhibiting water uptake and endosperm rupture (Finch-Savage and Leubner-Metzger, 2006). When favourable conditions are restored, abscisic acid levels decrease, with a concomitant increase of gibberellic acid (GA) to allow embryos to expand and break the seed covering layers (Manz et al., 2005). The endogenous levels of ABA and GA are regulated by different signalling pathways, and recent studies highlighted the crosstalk between light and hormonal pathways in the regulation of germination (Kim et al., 2008; Lau and Deng, 2010; de Wit et al., 2016). Phytochrome A (phyA) and phyB are photoreceptors which perceive Far Red (FR) and Red (R) light, respectively. Early during germination, phyB signalling involves a family of basic helix-loop-helix TFs, the PHYTOCHROME INTERACTING FACTORs (PIFs). After R or white light illumination, phyB translocates to the nucleus in its active Pfr conformation, where it binds and inhibits PIF1, also known as PIF3-LIKE 5 (PIL5), promoting light-induced germination (Lee et al., 2012). In the dark, or in low R/FR ratio light, when phyB is in the inactive, Pr cytosolic form, PIF1 is stabilized and represses germination. PIF1 promotes ABA biosynthesis and signalling, and represses GA signalling, inducing expression of genes such as *ABI3*, *ABI5, REPRESSOR OF GA1-3 (RGA), DOF AFFECTING GERMINATION 1 (DAG1)* (Oh et al., 2009). Interestingly, ABI3 protein also interacts with PIF1 to activate the expression of direct targets, such as *SOMNUS* (*SOM*), a key regulator of light-dependent seed germination acting on ABA and GA biosynthetic genes (Kim et al., 2008; Park et al., 2011).

In seeds initiating germination, *ABI3* expression is repressed. Perruc and colleagues (2007) reported that the chromatin-remodeling factor PICKLE negatively regulates *ABI3* by promoting silencing of its chromatin during seed germination. ABI3 activity is also controlled by alternative splicing of the corresponding precursor mRNA (pre-mRNA), with different splice forms predominating at different seed developmental stages. This process is regulated by splicing factor SUPPRESSOR OF ABI3-5 (SUA) through the splicing of a cryptic intron in ABI3 mRNA (Sugliani et al., 2010).

Alternative splicing occurs when the spliceosome differentially recognizes the splice sites. The selection of alternative 5’SS or 3’SS leads to an inclusion of different parts of an exon, whereas failure to recognize splicing sites causes intron retention in the mature mRNA. These alternative splice forms can produce proteins with altered domains and function (Staiger and Brown, 2013; Laloum et al., 2018; Nilsen and Brenton Graveley, 2010, Fu and Ares, 2014). In plants, this mechanism is highly induced in response to external stimuli. Recent studies reported an emerging link between splicing and ABA signalling (Zhu et al., 2017; Laloum et al., 2018). For example, the transcript encoding type 2C phosphatase HYPERSENSITIVE TO ABA 1 (HAB1), a negative regulator of ABA signalling, undergoes alternative splicing. In the presence of ABA, the last intron is retained, leading to a truncated protein. The two encoded proteins, *HAB1-1* and *HAB1-2*, play opposite roles by competing for interaction with OPEN STOMATA 1 (OST1) during germination, which then results in switching of the ABA signalling on and off (Wang et al., 2015). Likewise, SR45, a member of serine/arginine-rich proteins, an important class of essential splicing factors that influence splice site selection, regulates glucose signalling through downregulation of ABA pathway during seedling development (Carvalho et al., 2010). In addition, several splicing regulators were reported to influence ABA sensitivity, such as *SAD1*, *ABH1*, *SKB1, Sf1* (Xiong et al., 2001; Hugouvieux, et al. 2001; Zhang et al., 2011; Jang et al. 2014).

In this study we show that the splicing factor DNA-DAMAGE REPAIR/TOLERATION PROTEIN 111 (DRT111), previously characterized in the control of pre-mRNA splicing in light-regulated developmental processes (Xin et al., 2017), is involved in ABA response mechanisms. Manipulation of *DRT111* expression results in a modified sensitivity to ABA of stomatal movements and during seed germination. Accordingly, *DRT111* is highly expressed in stomata and seeds, and up-regulated upon long-term exposure to ABA. Moreover, *ABI3* alternative transcript quantification as well as analysis of double and triple mutants shows that *DRT111* controls splicing of ABI3 upstream of SUA. Transcriptome analysis in *drt111* dry seeds revealed extensive alteration in gene expression and splicing of genes involved in light and ABA-dependent control of germination. Consistently, we show that expression of the germination inhibitor *SOM* is induced in *drt111*. Taken together, our results suggest that integration of ABA and light quality stimuli for seed germination under appropriate conditions requires DRT111.

## Results

### *DRT111* expression is high in seeds and guard cells and is induced by long-term stress

Using RNA transcriptome data from potato cells adapted to gradually increasing concentrations of polyethylene glycol (PEG) we identified Arabidopsis orthologous genes, and functionally analysed their role in responses to ABA and osmotic stress (Ambrosone et al., 2015, 2017; Punzo et al., 2018). Following the same rationale, we focused on a DNA-damage-repair/toleration protein coding gene (EST617924, GenBank accession no. BQ510509, corresponding to PGSC0003DMT400054608, Potato genome sequencing consortium, 2011), up-regulated in adapted cells (Ambrosone et al., 2017). The protein deduced from PGSC0003DMT400054608 shared 64% sequence identity with Arabidopsis At1g30480, encoding a predicted splicing factor (also referred to as DRT111/ RSN2/SFPS, SPF45-related, Pang et al., 1993; Xin et al., 2017; Zhang et al., 2014, Supplemental Figure S1).

To verify the expression pattern in Arabidopsis, we mined public databases (eFP platform, Winter et al., 2007), showing that *DRT111* is ubiquitously expressed throughout development, with highest transcript abundance in dry seeds (Fig. 1A). In addition, histochemical analysis of stable Arabidopsis lines expressing β-glucuronidase (GUS) under the control of the *DRT111* promoter visualized GUS activity in cells surrounding the base of trichomes and guard cells (Fig. 1B).

**Figure 1.**
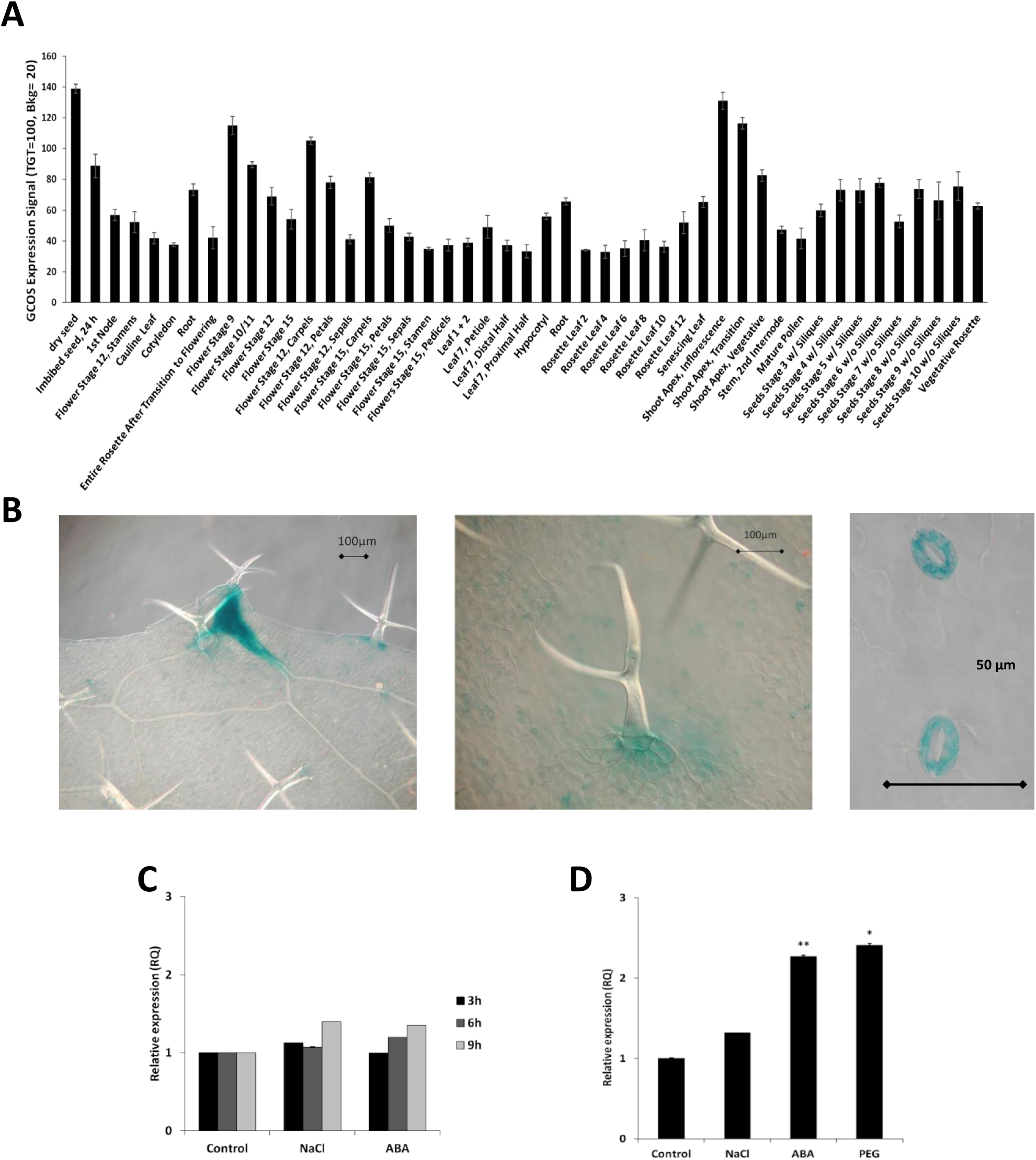
*DRT111* promoter activity and gene expression. (**A**) *DRT111* tissue-specific expression based on *Arabidopsis* microarray data in the eFP browser (http://bar.utoronto.ca). Data are normalized by the GCOS method, TGT value of 100. (**B**) Histochemical localization of GUS in leaves of transgenic *Arabidopsis* adult plants expressing the *GUS* reporter gene driven by *DRT111* promoter (DRT111promoter::GUS). Scale bars are shown. (**C**) Relative expression of *DRT111* in 7-day-old seedlings after 3, 6 and 9 h exposure to NaCl (120 mM) or ABA (50 μM) (**D**) Relative expression of *DRT111* in 7-day-old seedlings after 5 days exposure to NaCl (120 mM), ABA (10 μM) or polyethylene glycol (PEG; 35% W/V)). Data were normalized using RNA from untreated seedlings, and the elongation factor *EF1a* as endogenous control. Data reported are means ±SD of three biological replicates. The asterisks indicate significant differences compared to control condition according to Student’s t-test (*P ≤ 0.05, **P ≤ 0.01).

When assessing responsiveness of *DRT111* in seedlings exposed to short- or long-term treatments with NaCl, abscisic acid (ABA) or PEG, we detected a significant up-regulation of *DRT111* only after long-term treatments, while 3, 6 or 9h treatments did not result in major changes of *DRT111* transcript abundance. In particular, an increase higher than 2-fold was observed after 5-days treatments with ABA or PEG (Fig. 1C-D). Taken together, the results show that *DRT111* is highly expressed in seeds and guard cells, and that a higher steady state mRNA level is determined by long-term exposure to ABA or osmotic stress.

*DRT111/SFPS* encodes a nuclear-localized potential orthologue of human splicing factor *RBM17/SPF45* (Supplemental Figure S2, Xin et al., 2017), shown to interact with Splicing factor 1 (SF1), a protein involved in early pre-spliceosome assembly (Crisci et al., 2015; Hegele et al., 2012). Therefore, we verified if this interaction is conserved in Arabidopsis. Using the yeast two hybrid system, we tested different portions of SF1, and showed interaction between DRT111 and the C-terminal fragment (1398-2415) of SF1 (Fig 2A), while a fusion of the GAL4 binding domain (BD) with the SF1 N-terminal fragment (1-1396) resulted in auto-activation when co-transformed with the empty AD vector in yeast. We used a split reporter system to confirm the interaction *in planta*. A reconstituted YFP signal was detected in nuclear speckles, indicating that DRT111 forms a complex with SF1 and may thus act at early steps of the spliceosome machinery (Fig 2B).

**Figure 2.**
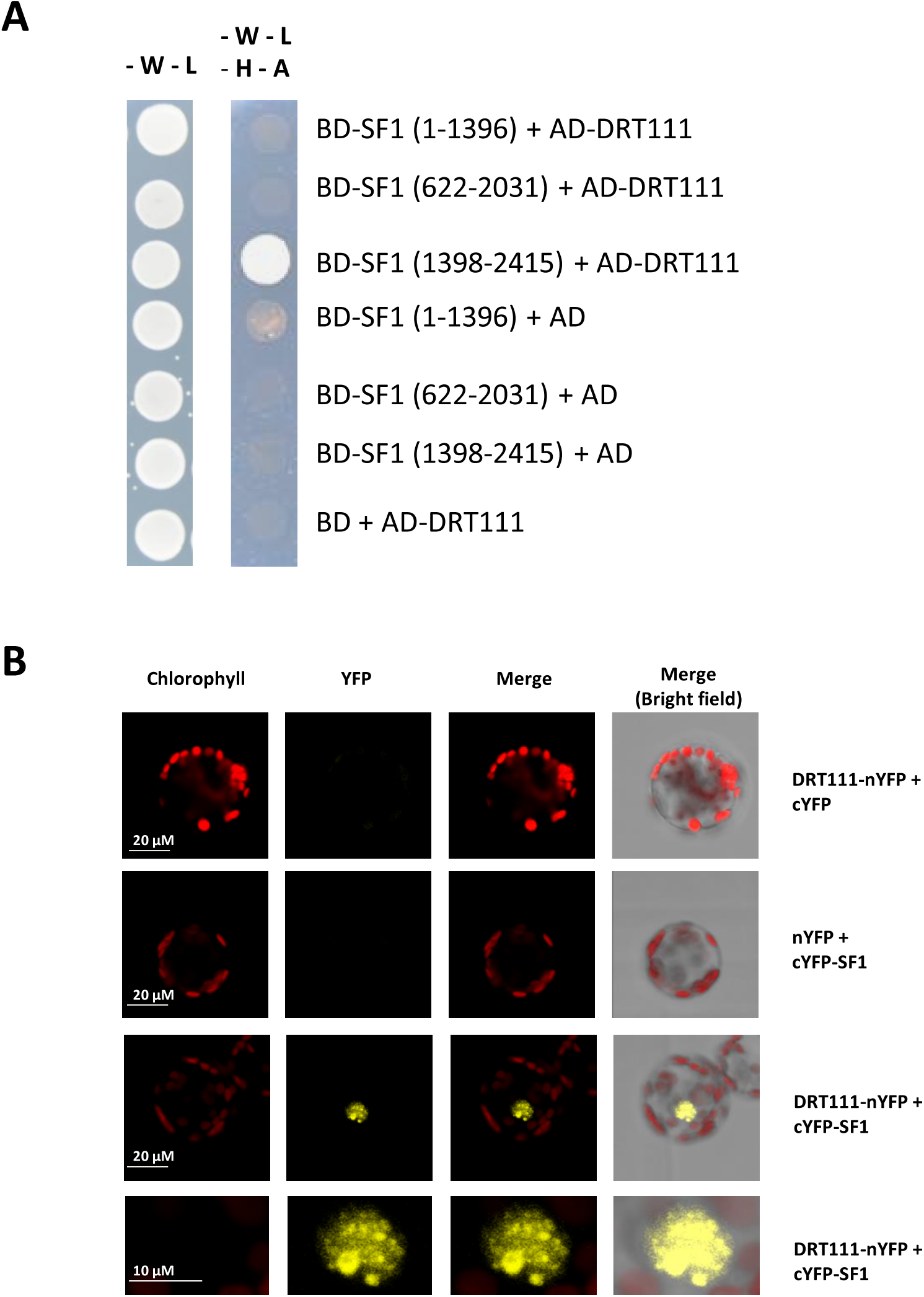
Interaction of DRT111 with Splicing Factor 1 (SF1). (**A**) Yeast two-hybrid assay. DRT111 in prey vector (pGADT7, AD domain) was co-transformed with the indicated fragments of SF1 cloned in the bait vector (pGBKT7, BD domain). The empty vectors pGADT7 and pGBKT7 were used as negative controls. Overnight grown yeast culture was dropped onto selective media. Pictures were taken after 3 days incubation at 30°C. (**B**) Bimolecular fluorescence complementation assay. *Nicotiana tabacum* leaf protoplasts were co-transformed with 20 μg each of plasmids encoding DRT111 fused with N-terminus of YFP (nYFP) and SF1 fused with the C-terminus of YFP (cYFP) . nYFP and cYFP empty vectors were used as negative controls. The cells were imaged by confocal microscopy 16 h later. For the interaction, zoom in images of the nucleus are shown in the last row. Chlorophyll autofluorescence, YFP fluorescence and merged images are shown. Scales bars are indicated.

### Altered *DRT111* expression affects plant growth and stomatal responsiveness to ABA

To analyse the involvement of DRT11 in ABA-related processes, insertion mutants were identified within the TAIR collection and transgenic plants over-expressing DRT111 were produced (Fig. S3A-C). Three over-expressing (OX) lines carrying homozygous, single-copy transgene insertions and expressing the transcript were selected (Fig. S3C). Phenotype observation in control conditions and after ABA treatment indicated that over-expression of *DRT111* caused a limited increase in primary root length (Fig. 3A), while lack of *DRT111* expression resulted in early flowering, as previously reported (Fig. S3D, Xin et al., 2017).

**Figure 3.**
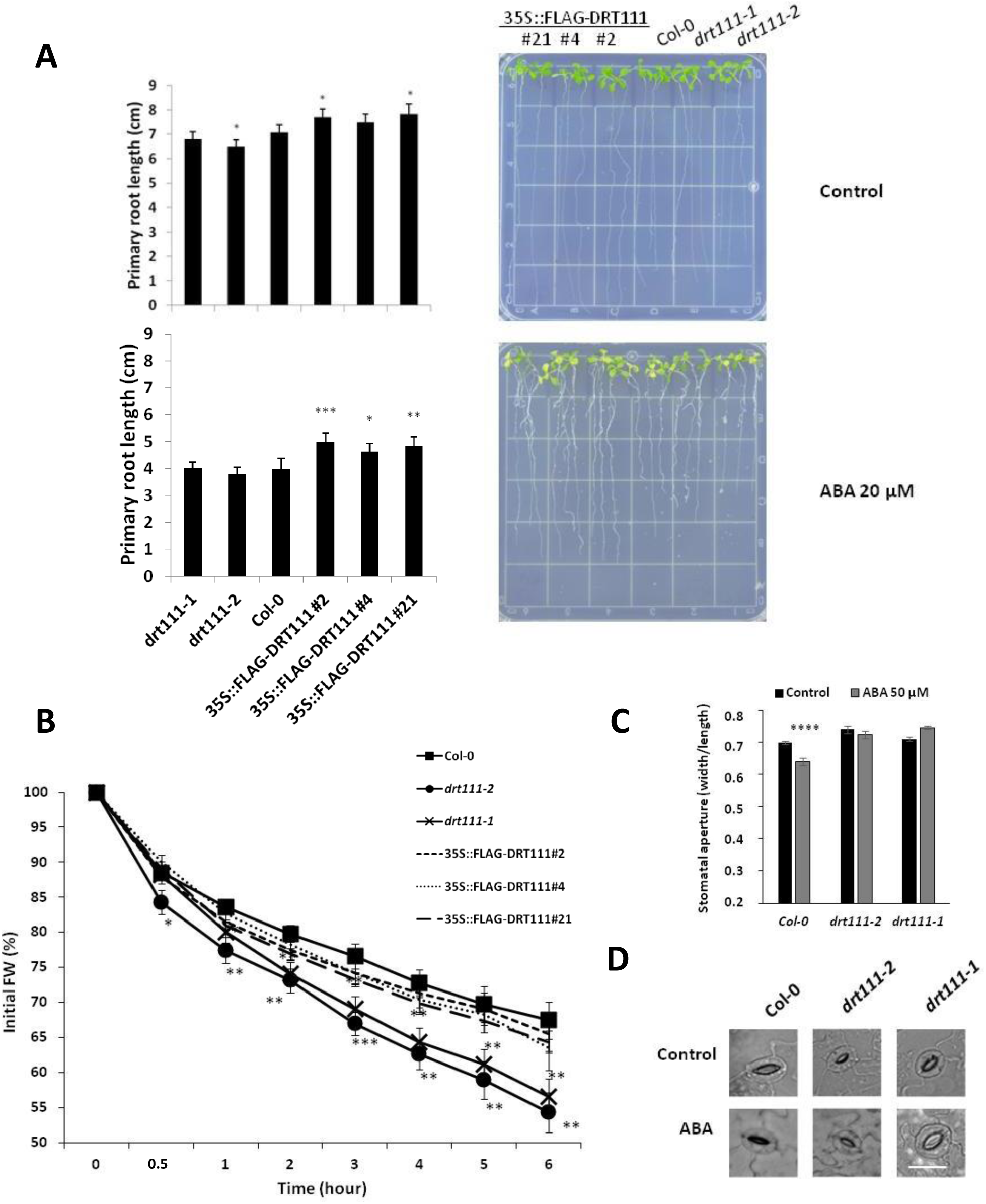
Phenotyping of knockout mutants *drt111-1* (GABI_351E09), *drt111-2* (SALK_001489) and *DRT111* over-expressing lines (35S::FLAG-DRT111 #2, 4, 21) (**A)** Primary root length of 10-day-old wild type (Col-0), DRT111 mutants and transgenic lines grown on GM medium (1% sucrose) or medium containing ABA 20 µM. (**B)** Water loss of detached leaves of *drt111* mutants, overexpressing lines (35S::FLAG-DRT111) and wild type (Col-0) plants. Data are averages ± SE of two independent experiments (n=5 for each line, per experiment) and reported as percentages of initial fresh weight at different time points (0.5 to 6 hours). The asterisks indicate significant differences compared with wild type (*P ≤ 0.05, **P ≤ 0.01, ***P ≤ 0.001, ****P ≤ 0.0001, Student’s t-test). (**C**) Stomatal aperture of *drt111-1* and *drt111-2* mutants and wild type plants in response to ABA. Leaf peels harvested from 2-week-old plants were incubated for 2 hours in SOS buffer under light and then treated with or without 50µM ABA for 2.5 hours. Asterisks indicate significant difference between sample with or without ABA (** P ≤ 0.01; Student’s t-test). **D**) Photographs of stomata of the indicated genotypes as reported in **C**. Scale bar: 25 µm.

As *DRT111* is highly expressed in guard cells (Fig. 1B), we evaluated the transpirational water loss and its relation to stomatal movements in plants with altered expression of *DRT111*. First, we measured the fresh weight reduction of detached leaves during six hours (Verslues et al., 2006). Whereas the OX and wild-type plants showed similar trends, we observed a significant increase in the transpirational water loss in knockout plants, with a loss of 44% and 46% of their initial fresh weight for *drt111-1* and *drt111-2*, respectively compared to 33% of Col-0 after 6 hours (Figure 3B). Thus, we analyzed the stomatal movements in *dtr111* plants compared to wild type after treatments with abscisic acid, which plays a central role in stomatal closure (Figure 3C-D). Consistent with the water loss analysis, significant differences in stomatal pore size were observed between genotypes after 2.5h ABA treatments (50µM). While the ABA-induced stomatal closure was observed in Col-0 leaves (ratio 0.64 width/length of pore, corresponding to 8.5 % reduction of the pore size), stomata of the mutants did not respond to the ABA treatment (*drt111-1*) or had a strikingly reduced response (*drt111-2*, ratio 0.72 width/length of pore, corresponding to 2.2% reduction compared to untreated stomata), suggesting that stomatal responsiveness to ABA is impaired in *drt111* mutants causing a significant water loss over time.

### drt111 seeds are hypersensitive to ABA during germination

Since *DRT111* is an ABA-responsive gene highly expressed in seeds (Fig.1A), we analysed seed germination of mutants and over-expressing lines in presence of ABA. As indicated in Fig. 4, seeds collected from *drt111-1* and *drt111-2* were hypersensitive to ABA in terms of radicle emergence and cotyledon expansion, while OX lines displayed an increased seed germination in presence of ABA compared to wild type. In particular, after 3 days 77% of Col-0 seeds were germinated on 0.5 µM ABA, compared to 96, 93 and 93% of FLAG-DRT111 #2, #4 and #21, respectively and 53, 57% in *drt111-1* and *drt111-2* (Fig. 4A-C). An ABA response curve of 12 months after-ripened seeds indicated that the hypersensitivity at high concentrations of ABA of the *drt111* mutants was retained after long periods of dry storage (Fig. 4D). Complementation experiments of *drt111* with *DRT111* driven by the endogenous promoter, indicate that the hypersensitivity to ABA of knockout *drt111-1* and *drt111-2* can be reverted by introducing a functional *DRT111* copy, thus confirming that mutant phenotype is caused by lack of functional *DRT111* (Fig. 4E).

**Figure 4.**
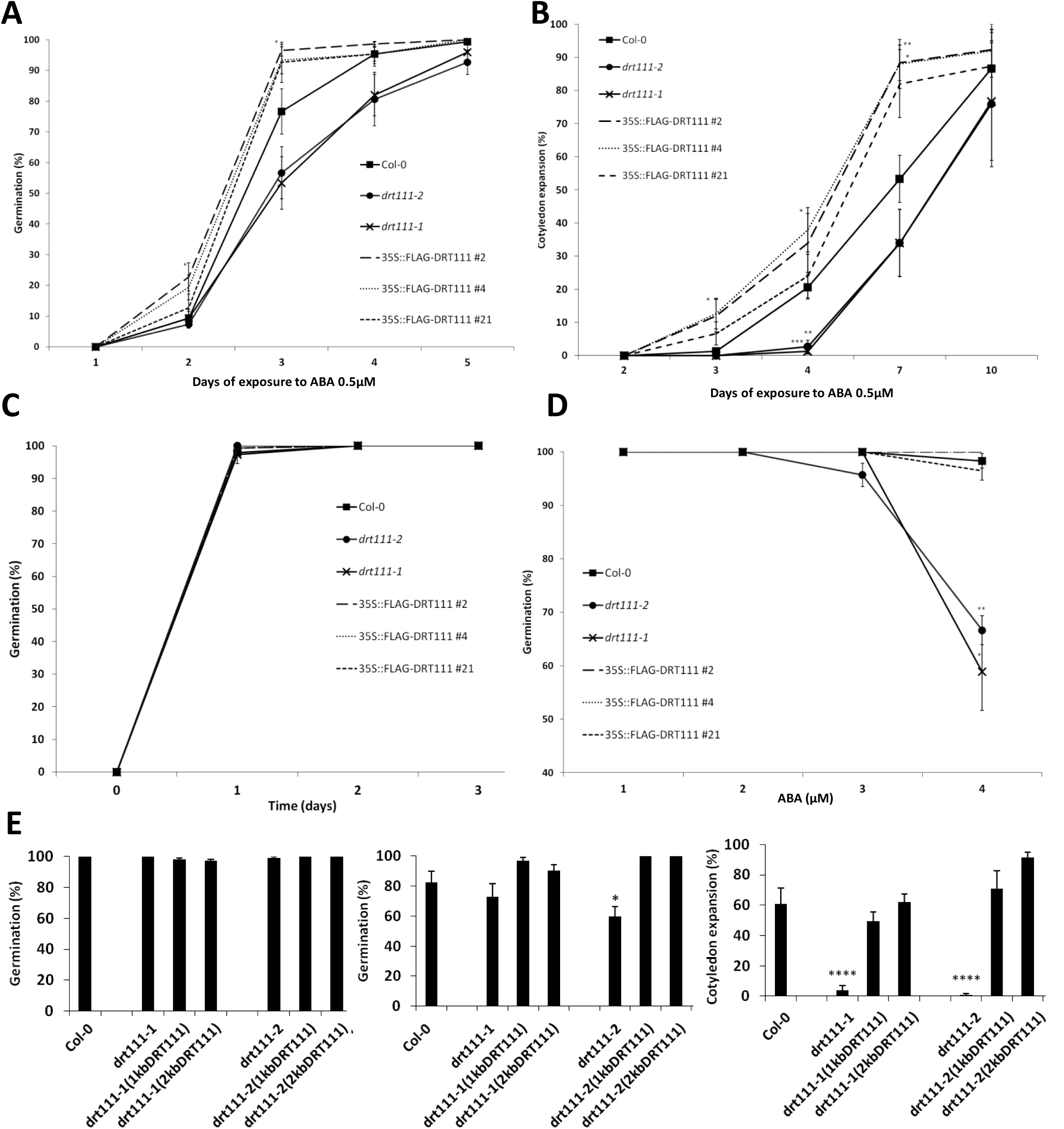
Germination analysis of *drt111* mutants (*drt111-1*, *drt111-2*), *DRT111* over-expressing lines (35S:FLAG-DRT111 #2, #4, #21), and wild-type (Col-0) and complementation of *drt1111* mutants with pDRT111:DRT111-FLAG (**A-C**) Germination percentage of 10 days after-ripened seeds scored in terms of radicle emergence (**A**) and cotyledon expansion (**B**) in presence of 0.5 µM ABA and control media (**C**). Data are means ±SD (n=150) of five biological replicates. **D)** Germination percentage of 1 year after-ripened seeds scored in terms of radicle emergence after 3 day in the presence of different concentration of ABA. **E**) Germination analysis of *drt111* mutants transformed with *DRT111* genomic fragment including 1kb or 2kb upstream of the translation start site (1KbDRT111; 2kbDRT111) in pDRT111:DRT111-FLAG constructs. Germination reported as percentage in terms of radicle emergence in control condition (left) and in the presence of 0.5 µM ABA (middle) and in terms of cotyledon expansion in presence of 0.5 µM ABA (right). In all germination tests, seeds were stratified for 2 d before incubation at 24°C. The asterisks indicate significant differences compared to Col-0 according to Student’s t-test (*P ≤ 0.05, **P ≤ 0.01, ***P ≤ 0.001, ****P ≤ 0.0001).

### DRT111 regulates gene expression and mRNA splicing

To further characterize the role of *DRT111* in seed germination, we examined the transcriptome of *drt111* dry seeds. RNA-seq analysis highlighted a major role of *DRT111* in the regulation of gene expression with over 3000 differentially expressed genes (ǀlog2(fold-change)ǀ> 0.21, FDR <0.05), equally distributed among down-(1941 genes) and up-regulated (1834) genes (Supplemental Dataset 1). Validation of a subset of genes by qRT-PCR showed high correlation with the fold change detected by RNA-seq (Supplemental Figure S4C-D).

Consistent with the observed phenotype, gene ontology (GO) enriched categories included seed related processes (seed germination, embryo development ending in seed dormancy, post-embryonic development), response to abiotic stress (response to salt stress, response to cold, response to water deprivation, response to heat, regulation of stomatal movement, hyperosmotic salinity response, response to osmotic stress), in the ABA signalling pathway (response to abscisic acid, abscisic acid-activated signaling pathway,) as well as the processing of mRNAs (mRNA processing, RNA splicing) (Supplemental Dataset S1, Figure S4A-B).

Among the genes differentially expressed in *drt111-2*, components of the light perception/signalling cascade were present, including Phytochromes (Phy) and PHYTOCHROME INTERACTING FACTORs (PIF), some of which up-regulated (*PhyA*, *PIF1/PIL5* and *PIF6/PIL2*) and others down-regulated (*PhyE*, *PhyD* and *PIF7*) (Supplemental Dataset S1).

Significantly upregulated genes in *drt111-2* included several members of the homeodomain leucine zipper class I TF (*ATHB-1*, *ATHB-5*, *ATHB-7*, *ATHB-12*), which regulate abiotic stress responses.

To investigate the impact of lack of *DRT111* on pre-mRNA splicing, we explored differences in splicing events between *drt111-2* and Col-0. Using the MATS (Multivariate Analysis of Transcript Splicing) software, we analyzed all major types of splicing events, such as exon skipping (ES), alternative 5’ or 3’ splice site (A5SS; A3SS), mutually exclusive exon (MXE) and intron retention (IR). All the analyzed events were affected in *drt111-2* seeds. We identified a total of 611 differential splicing events, corresponding to 485 genes between mutant and wild type. Among these, A3SS and IR were the most rapresented categories, with 161 and 258 events respectively (Fig 5A-B, Supplementl Dataset S2). Interestingly, gene ontology enrichment analysis (GOEA) showed that categories related to germination mechanisms (such as response to abscisic acid, positive regulation of seed germination, abscisic acid biosynthetic process, maintenance of seed dormancy by absisic acid, regulation of seed germination, embryo sac egg cell differentiation) or to mRNA metabolism (such as mRNA splicing via spliceosome, mRNA processing, RNA splicing, mRNA stabilization) were significantly enriched among the IR and A3SS defects in *drt111-2*. (Supplemental Dataset S3), suggesting that DRT11 may control the splicing of specific mRNAs in seeds. We validated the splicing events identified through RNA-seq and reported as reads mapped in gene regions in Col-0 and *drt111-2* mutant (Fig 5C, E, G and I) by qRT-PCR analysis (Fig 5D, F, H and J).

**Figure 5.**
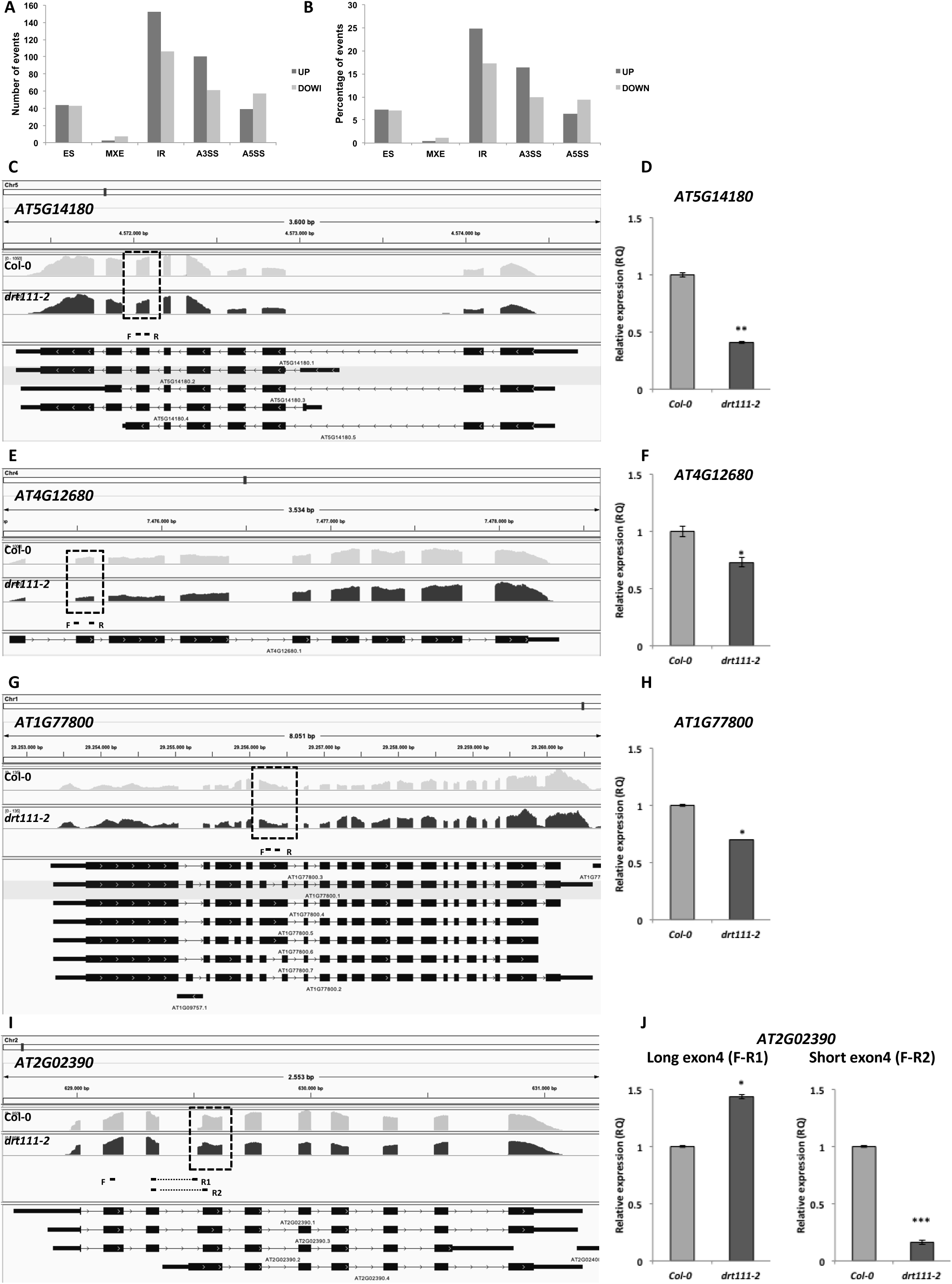
Alternative splicing (AS) events altered in *drt111-2*. **A**) Number of different AS events UP- (a greater prevalence of AS event in the mutant *vs.* Col-0) and DOWN- (a lower prevalence of AS event in the mutant *vs.* Col-0) regulated in *drt111-2* dry seeds. **B**) Percentage of splicing events significantly UP- and DOWN-regulated in *drt111-2* with respect to the total AS events defective in *drt111-2*. ES: Exon skipping; MXE: Mutually exclusive exon; IR: Intron retaining; A3SS: Alternative 3’ splice site; A5SS: Alternative 5’ splice site. (**C-J**) Validation of RNA-seq data by qRT-PCR. (**C, E, G** and **I**) Representation of AS differences between Col-0 and *drt111-2* detected by RNA-seq using Integrative Genomics Viewer. Dashed box indicates the position of alternative splicing events: ES (*AT5G14180*, *AT4G12680*), IR (*AT1G77800*) and A3SS in (*AT2G02390*). Primers used for qRT-PCR are shown. (**D, F, H** and **J**) Validation by qRT-PCR. The elongation factor EF1α was used as endogenous control.

### DRT111 regulates ABI3 splicing

We have shown that *drt111* mutants are hypersensitive to ABA in the germination process (Fig. 4). One of the key players determining sensitivity to ABA at the seed stage, and whose activity is regulated by alternative splicing, is the transcription factor ABI3 (Sugliani et al., 2010). The *ABI3* locus gives rise to two alternative transcripts, *ABI3-α* and *ABI3-β* which differ by the presence or cleavage of a cryptic intron, respectively. *ABI3-α* produces a full length, functional protein and is highly expressed during seed development, while *ABI3-β,* encodes a truncated protein lacking two of the four ABI3 conserved domains, and accumulates at the end of seed maturation (Sugliani et al., 2010).

Although splicing of *ABI3* was not identified as affected in *drt111-2* through RNAseq, comparison of the DEGs with a list of 98 *ABI3* targets (Monke et al., 2012) showed that 51 of these genes were deregulated in *drt111-2* (Supplemental Table S1), suggesting that *ABI3* might be a target of *DRT111*.

Therefore, we used qRT-PCR to quantify the amount of *ABI3-α* and *ABI3-β* in *drt111-2* dry and imbibed seeds compared to wild type using primers described by Sugliani and colleagues (2010). Although the level of *ABI3-α* is similar in dry seeds, accumulation of *ABI3-β* is significantly higher in *drt111-2* than Col-0; in addition, in imbibed seeds, both transcripts were upregulated in *drt111-2*, with *ABI3-β* showing a 4-fold induction compared to Col-0 (Fig. 6A), demonstrating a defective regulation of *ABI3* splicing in plants lacking *DRT111*.

**Figure 6.**
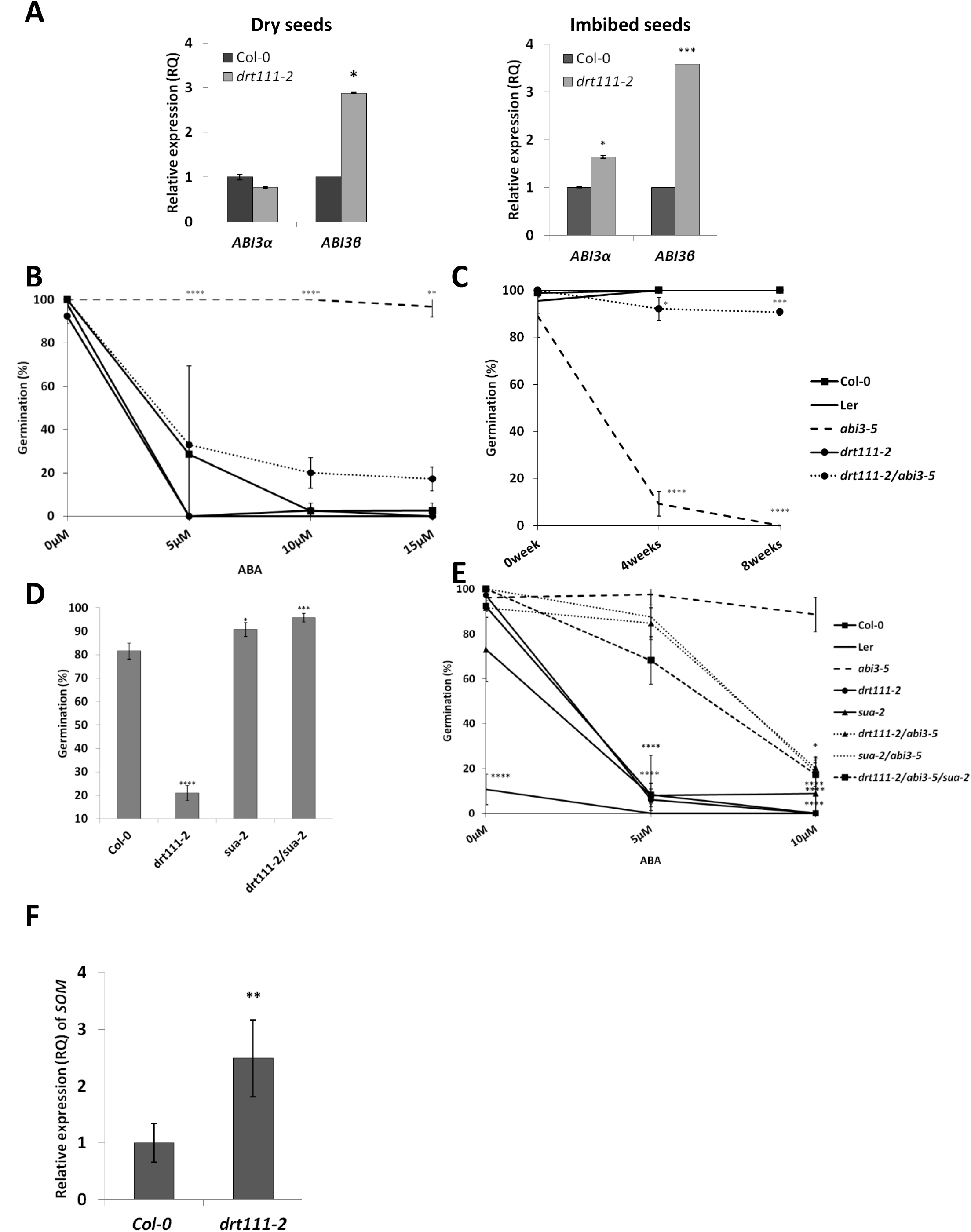
Germination analysis and relative expression of *ABI3* and *SOMNUS* in *drt111-2* seeds**. A)** Expression of *ABI3* splicing variants (ABI3-α, ABI3-β) in Col-0 and *drt111-2* dry seeds (left) or imbibed seeds (right). **B**) Germination of freshly harvested seeds sown on media containing 0, 5, 10 or 15µM ABA. **C**) Germination of seeds sown on GM medium after different periods of dry storage: 0, 4 or 8 weeks after harvest. (**B,C**) Data were collected after 3 days and reported as means of three biological replicates (±SD). **D**) Germination of 14 d after-ripened seeds sown on medium containing ABA 0.5 µM. Data were collected after 3 days and reported as means of two independent experiments (±SE). **E**) Germination of freshly harvested seeds sown on medium containing different concentrations of ABA (0; 5; 10µM). Data were collected after 7 days and reported as means of three biological replicates (±SD). **F**) Expression of *SOMNUS* in Col-0 and *drt111-2* imbibed seeds. In all germination tests, seeds were stratified for 2 d before incubation at 24°C. The asterisks indicate significant differences compared with wild type or *abi3-5* (**E**) according to Student’s t-test (*P ≤ 0.05, **P ≤ 0.01, ***P ≤ 0.001, ****P ≤ 0.0001).

To confirm this observation genetically, we took advantage of the *abi3-5* mutant allele (Ooms et al., 1993). Due to a frameshift mutation, the *abi3-5-α* transcript contains a premature stop codon, while *abi3-5-β* encodes a functional ABI3 protein, therefore an increase in accumulation of *abi3-5-β* results in a higher amount of functional ABI3 (Sugliani et al. 2010). Thus, an increased efficiency in splicing of the cryptic intron is expected to alleviate *abi3-5* phenotypes, including ABA insensitivity during germination and reduced seed longevity (Bies-Etheve et al., 1999).

We thus produced double mutants *drt111-2/abi3-5* to verify reversion of the *abi3-5* phenotypes. Germination tests showed increased sensitivity to ABA and improved longevity of *drt111-2/abi3-5* as compared to *abi3-5*. In the presence of 10µM ABA, 20% of *drt111-2/abi3-5* seeds were able to germinate compared to 100% in *abi3-5* (Fig 6B). Also the severe reduction of longevity observed in *abi3-5* seeds was restored in *drt111-2/abi3-5*, with 90% of seeds germinated 8 weeks after harvest (Fig 6C), compared to 0% *abi3-5*. Altogether, these results show that mutations in *DRT111* rescue the *abi3-5* mutation, with knock-out mutations showing a higher efficiency of phenotype reversion.

Since the alternative splicing of *ABI3* is also controlled by the splicing factor SUPPRESSOR OF ABI3-5 (SUA, Sugliani et al., 2010), we verified genetic interaction between *DRT111* and *SUA* by the analysis of the double mutants *drt111-2/sua-2*. As shown in Figure 7, seed germination on 0.5 µM ABA of *drt111-2/sua-2* was 96%, compared to 20% of *drt111-2* and 91% of *sua-2.* Thus, these results indicate that *SUA* is epistatic to *DRT111* and that *DRT111* acts upstream of *SUA* (Figure 6D). Accordingly, germination of *drt111-2/sua-2/abi3-5* triple mutant did not show additive effects compared to *drt111-2/abi3-5* or *sua-2/abi3-5* double mutants. Germination of the triple mutant on higher ABA concentrations largely resembled that of double mutants, with 17% *drt111-2/sua-2/abi3-5* seeds germinated on 10µM ABA compared to 20% *drt111-2/abi3-5* and 18% *sua-2/abi3-5*, indicating that control of *ABI3* splicing by *DRT111* and *SUA* is exerted through the same pathway (Figure 6E).

### Expression of SOMNUS is affected in drt111

In imbibed seeds, ABI3 and PIL5/PIF1 collaboratively activate the expression of the germination inhibitor *SOMNUS* (*SOM,* Park et al., 2011), whereas R or white light repress it through the action of PhyB. Since *DRT111* controls splicing of ABI3, and is epistatic to *PIFs* (Xin et al., 2017), we verified expression of *SOM* in imbibed seeds of *drt111-2.* As shown in Fig. 6F, expression of *SOMNUS* was 2.49-fold higher in *drt111-2* compared to wild-type Col-0, indicating that a higher expression of *SOM* might contribute to the observed ABA hypersensitivity in *drt111* seeds. Consistently, *pil5/pif1* and the quadruple *pif1/pif3/pif4/pif5* mutant (*pifq*) are insensitive to ABA in seed germination (Supplemental Figure S5 B-C; Oh et al., 2009; Lee et al., 2012). Finally, we also find that the *phyB* mutant is hypersensitive to ABA in the germination process (Fig S5A). Again, this could be partly due to the lack of negative regulation of SOM acting as a positive regulator of ABA biosynthesis (Kim et al., 2008).

## Discussion

Alternative splicing and its regulation are involved in several adaptation processes in response to environmental stimuli and stresses (Laloum et al., 2018). Here, we have shown that the Arabidopsis *DRT111* gene, encoding a protein orthologous to the human splicing factor SPF45 (Xin et al., 2017), is highly expressed in dry seeds, stomata and in seedlings experiencing long-term osmotic stress. Functional studies in Arabidopsis established that DRT111 controls stomatal movements and seed germination in response to ABA.

The human splicing factor SPF45 forms a complex with SF1 and U2AF^65^ for the selection of alternative 3′ splice sites (Lallena et al. 2002). During early spliceosome assembly, the U2AF^65^ interact with U2AF^35^ and SF1 to promote U2snRNP detection of the pre-mRNA 3’ splice site (Park et al., 2017).

Here we have shown that DRT111 physically interacts with SF1, while, in a previous independent study, interaction and colocalization with U2AF^65^ was reported (Xin et al., 2017). Based on homology with yeast and metazoan proteins, plant SF1 may be involved in recognition of intron branching point and assist in 3’ splice site selection (Jiang et al., 2014; Lee et al., 2017). Together with the observation that the highest number of the observed splicing defects concerned IR and A3SS, the protein interactions suggest that DRT111 is also involved in the early steps of spliceosome formation, which concern intron branch point recognition and 3’ splicing site selection by U2AF and SF1. However the mode of participation of DRT111 (e.g. promotion and/or suppression) in this complex remains to be established.

A growing body of evidence indicates that in plants components of the pre-mRNA splicing machinery modulate responses to ABA and abiotic/biotic stresses (Xiong et al., 2001; Cui et al., 2014; Carrasco-Lopez et al., 2017). Arabidopsis *sf1* mutants show several developmental defects, including dwarfism, early flowering and hypersensitivity to ABA at seed germination stage (Jiang et al., 2014).

Previously, *DRT111* and SUPPRESSOR OF ABI3-5 (SUA) were identified in a suppressor screen of *snc4-1d*, mutated in a receptor like kinase involved in bacterial pathogen resistance (Zhang et al., 2014). A similar pattern of intron retention in *SNC4* and *CERK1* was reported in both *sua* and *drt111* plants, thus suggesting that *SUA* and *DRT111* are both required for the splicing of at least these two genes (Zhang et al., 2014).

Here, we have shown that DRT111 knock-out and over-expressing plants are impaired in ABA seed germination responses, in analogy to *sua* mutants (Sugliani et al., 2010). In particular, SUA controls the activity of ABI3 by suppressing the splicing of an *ABI3* cryptic intron to reduce the levels of functional ABI3 in mature seeds (Sugliani et al., 2010). Because the *ABI3* cryptic intron is part of a protein-coding exon, it was subsequently classified as an exitron, an alternatively spliced internal region of a protein-coding exon (Marquez et al., 2015). Exitron splicing (EIS) is suggested to be a mechanism to increase plant proteome diversity in specific developmental stages or stress conditions, to affect protein functionality by modifying intracellular localization, presence of protein domains and post-translational modification sites, such as phosphorylation, sumoylation, ubiquitylation (Marquez et al., 2015). Based on EIS patterns in *sua* mutants, and presence of RBM5/SUA predicted binding sites enrichment in exitrons, SUA appears to have a general role in preventing exitron splicing (Marquez et al., 2015).

Here, we have shown that DRT111, similarly to SUA, suppresses splicing of *ABI3;* accordingly, known ABI3 targets (Monke et al., 2012) were found differentially expressed in *drt111* compared to wild type. Interestingly, *sua* mutants in Columbia background are insensitive to ABA in seed germination (Sugliani et al., 2010), whereas *DRT111* knock-out causes ABA hypersensitivity. The phenotype in *drt111* may be explained by the observed increase in total *ABI3* amount, determined in imbibed seeds by an increase of both the α and the β transcripts. In particular, a 4-fold accumulation of ABI3-β, corresponding to the transcript in which the exitron is spliced out, accounts for most of *ABI3* transcript in *drt111-2*. Therefore, the different ratio between the *ABI3-α* and *ABI3-β* transcripts, and their products thereof, may be important to define seed ABA sensitivity.

Both *sua/abi3-5* or *drt111-2/abi3-5* in Columbia background partially rescue seed developmental and ABA sensitivity defects of *abi3-5.* Thus, similarly to mammalian systems, SUA and DRT111 may control splicing of the same substrates with different timing. Further analyses will verify if DRT111 also controls exitron splicing mechanism.

DRT111/SFPS was recently shown to regulate development in response to light through interaction with phyB and REDUCED RED-LIGHT RESPONSES IN CRY1CRY2 BACKGROUND 1 (RRC1, Xin et al., 2017, Xin et al., 2019). In vegetative tissues, DRT111 regulates pre-mRNA splicing of genes involved in light signaling and the circadian clock and acts upstream of PHYTOCHROME INTERACTING FACTORS (PIFs) transcription factors, a major class of phyB targets (Xin et al., 2017). Interestingly, we observed a differential expression of *PIF1/PIL5*, *PIF6/PIL2* and *PIF7* in dry seeds of *drt111-2*: in particular *PIF1/PIL5* and *PIF6/PIL2* were upregulated and *PIF7* was slightly down-regulated.

PIF1 inhibits GA signalling by promoting expression of DELLA repressors and, indirectly, by reducing GA levels (Oh et al., 2007; Paik et al., 2017). Indeed, up-regulation in *drt111-2* seeds observed for GIBERELLIC ACID INSENSITIVE (GAI) and *RGA-LIKE2 RGL2,* could also be dependent on an increased *PIF1* activity or expression (Lee et al., 2012).

In the dark, or in response to low R/FR ratio light, PIF1 inhibits seed germination through activation of hormone-dependent, germination-inhibiting mechanisms, including the induction of ABA biosynthesis and signalling genes (Oh et al., 2009). This process is partly regulated by the action of SOM (Kim et al., 2008; Park et al, 2011) which in turn regulates MOTHER-OF-FT-AND-TFL1 (Vaistij et al. 2018). This may be achieved through induction of expression and interaction with ABI3 and ABI5, which may assist PIF1 in target site selection and activation of transcription (Kim et al., 2008; Park et al., 2011; Kim et al., 2016). Here we have shown that expression of *SOM* is upregulated compared to wild-type in *drt111-2* imbibed seeds. Thus, regulation of *SOM* appears to be a major point of convergence of light and hormonal stimuli and *DRT111* may be involved in this signal integration by exerting a regulatory function on both *ABI3* and *PIF1*.

Since *phyB-9* seeds are hypersensitive to ABA and it has been previously shown that phyB plants maintain open stomata under stress conditions, similarly to what we observed in *drt111* mutants (Gonzalez et al., 2012), we cannot exclude that the light perception by phyB is involved in DRT111-dependent splicing events.

Finally, the transcriptomic analysis identified several genes whose expression/splicing is affected in *drt111-2,* therefore, several other key factors may contribute to the observed ABA hypersensitivity in *drt111*. Among them, genes highly expressed in *drt111-2* included several members of the homeodomain leucine zipper class I TF (ATHB-1, ATHB-5, ATHB-7, ATHB-12), which have been largely studied for their role as regulators of abiotic stress responses. *ATHB-7* and *ATHB-12* are induced by water stress and ABA and control expression of several members of clade A PP2Cs, and are therefore considered negative regulators of ABA and stress responses (Arce et al., 2011; Valdes et al., 2012; Sessa et al., 2018). On the contrary, *ATHB-5* whose expression is positively regulated by ABI1, ABI3, ABI5, is considered a positive regulator of ABA signaling since enhanced levels of ATHB-5 result in elevated ABA responses (Johannesson et al., 2003). *ATHB-1*, in particular, was shown to be regulated at the expression level by PIF1/PIL5 and regulates hypocotyl growth in short days (Capella et al., 2015). Future work will analyse the molecular details of the regulation operated by DRT111 on its targets.

AS defects in *drt111* concerned predominantly IR and A3SS. Other splicing effectors and regulators affecting stress responses regulate these two AS classes. An increased splicing efficiency of IR prone introns was shown to be important for acclimation to drought stress and splicing regulator HIN1 is involved in this process (Chong et al., 2019). Similarly, SAD1 splicing factor increased A3SS usage under salt stress conditions (Xing et al., 2015). How DRT111 and components/regulators of the spliceosome, including SUA, SF1, U2AF^65^ associate/compete to determine the splicing of specific transcripts will be important to establish the contribution of this layer of regulation in defining the proteome during ABA and stress responses.

In conclusion, ours and previous evidence shows that DRT111 constitutes a point of integration of light and ABA-dependent signaling by controlling expression and splicing of key factors such as *ABI3* and *PIF* transcription factors.

## Experimental Procedures

### Plant materials, growth conditions and germination assays

The Columbia (Col-0) and Landsberg (Ler) ecotypes were used as wild-type. The *drt111* T-DNA insertion mutants: *drt1111-1* (GABI_351E09), *drt111-2* (SALK_001489) were obtained from the Nottingham Arabidopsis Stock Centre (NASC). *sua-2* and *sua-2/abi3-5* were kindly donated by Professor Wim Soppe (Max Planck Institute for Plant Breeding Research, Germany; present address Rijk Zwaan, Netherlands). *abi3-5* was donated by Dr. Lucio Conti (Department of Biosciences, University of Milan, Italy). *Arabidopsis thaliana* plants were grown on soil in a growth chamber (14 h light /10 h dark) at 24°C. For germination tests, seeds harvested the same day from plants grown in parallel and stored for the same time were compared. Freshly harvested seeds or dry stored (after-ripened) for different times as indicated in figure legends were used. Seeds were sown on GM medium (1X MS salts, 0.5% sucrose, pH 5.7) or medium containing different concentrations of ABA (0.5 µM, 2 µM, 5 µM, 10 µM). After stratification treatment at 4°C for 2 days, seeds were transferred to a growth chamber (16 h light / 8 h dark) at 24°C. Germination percentage was evaluated in terms of radicle emergence or fully expanded cotyledons. Gene expression analysis was carried out using 7-day-old seedlings grown on GM plates and then transferred to GM or GM containing NaCl (120 mM) and ABA (50 µM) for 3, 6 and 9 h, or NaCl (120 mM), ABA (10 µM) or PEG (35% w/v) for 2 and 5 days.

### RNA extraction, cDNA synthesis and qRT-PCR

Total RNA was isolated from 100 mg of seedlings using RNeasy Plant Mini Kit (Qiagen, Hilden, Germany) according to manufacturer’s instructions. For RNA deep sequencing and qRT-PCR, total RNA was extracted from 100mg of dry seeds or imbibed seeds (in H_2_O, 24h in dark, 4°C) using method reported in Oñate-Sánchez and Vicente-Carbajosa (2008). cDNA was synthesized using the QuantiTect Reverse Transcription Kit (Qiagen, Hilden, Germany), starting from 1µg of DNase-treated RNA. For qRT-PCR, 4.5 µl of diluted (1:20) cDNA was used for each reaction, with 6.25 µl of 1X Platinum SYBR Green qPCR SuperMix (Life Technologies, Carlsbad, CA, USA) and 1.75 µl of primer mix (5 µM). PCR was performed using ABI 7900 HT (Applied Biosystems, Foster City, CA, USA). Cycling conditions were: 10 min at 95°C, followed by 40 cycles of 95°C for 15s and 60°C for 1 min. Three biological replicates, each with three technical replicates were tested. The relative quantification of gene expression was calculated based on the 2^-ΔΔCt^ method (Livak and Schmittgen, 2001). The elongation Factor *EF1α* was used as endogenous reference gene and RNA isolated from control plants as calibrator sample. Primers used are listed in Supplemental Table S2.

### Generation of DRT111 transgenic plants

Transgenic Arabidopsis plants were produced using binary vectors obtained by Gateway technology (Life Technologies, Carlsbad, CA, USA). To study promoter activity, the sequence of *DRT111* promoter (2kb upstream of the start codon) was amplified from genomic DNA of Col-0 plants. To permit both N-terminus than C-terminus fusion with tags, the coding sequence of *DRT111* was amplified with or without STOP codon. For the complementation of *drt111* mutants, the genomic fragment of *DRT111* including the upstream 1kb or 2kb region were amplified. Primers used are listed in Supplemental Table S2. PCR amplifications were performed using Phusion DNA polymerase (Thermo scientific, Waltham, MA, USA). The amplicones were cloned into pDONR207 (Life Technologies, Carlsbad, CA, USA) using BP clonase (Life Technologies, Carlsbad, CA, USA) to obtain entry vectors.

Recombination with destination vectors was perfomed using LR clonase (Life Technologies, Carlsbad, CA, USA). pMDC164 (Curtis and Grossniklaus, 2003) was used for promoter studies, pGWB411 and pGWB412 (Nakagawa et al., 2007) to produce FLAG-tagged over-expressing plants, pEG302 (Earley et al., 2006) for mutant complementation. The resulting recombinant binary vectors were then introduced into the *Agrobacterium tumefaciens* GV3101 strain, which was then used to transform Col-0 plants or *drt111* mutants using the floral dip method (Clough and Bent, 1998).

### GUS assay

Histochemical analysis of GUS activity was performed as described previously (Batelli et al., 2012). The tissues from transgenic Arabidpsis plants transformend with *DRT111promoter*::GUS construct were washed in 70% ethanol and cleared with chloralhydrate/glycerol solution. Samples were analysed and photographed under an Axioskop 2 plus microscope (Zeiss) equipped with a Nikon Coolpix 990 camera.

### Stomatal measurements

Detached leaves from 4-week-old plants were used for stomatal measurements. For stomatal aperture assay, epidermal peels were floated in SOS solution (20 mM KCl, 1 mM CaCl2, and 5 mM MES-KOH pH 6.15) for 2.5h at light to induce stomatal opening. Then the buffer was replaced with fresh SOS containing 50µM of ABA or fresh SOS without ABA and incubated at light for 2.5h. 100 stomata were randomly observed using a Leica DMR microscope. The widths and lengths of stomata pores were measured using Image J software.

### Yeast two-hybrid assay

For the yeast two hybrid assay, the coding sequence of *DRT111* was cloned into the BamH1 and XhoI restriction sites of pGADT7 vector (Clontech, Mountain View, CA, USA) and the cDNA fragments of *SF1* were cloned into the SmaI and SalI sites of pGBKT7 (Clontech, Mountain View, CA, USA) using primers listed in Supplemental Table S2. To evaluate the interaction between DRT11 and the different SF1 fragments, the obtained constructs were co-transformed into *S.cerevisiae* AH109 strain using the LiAc-mediated transformation method (Bai and Elledge, 1996) and plated on SD medium (7.5 g/L Yeast Nitrogen Base, 0.75 g/L amino acid mix, 20 g L/1 glucose, pH 5.8) lacking Leu and Trp. Yeast cultures were grown overnight and an equal amount was dropped on SD lacking Leu and Trp medium to guarantee the presence of both vectors, and onto SD medium lacking Leu, Trp, Ade and His to verify the protein-protein interaction (Ruggiero et al., 2019). Empty vectors pGBKT7 and pGADT7 were used as negative controls.

### Bimolecular fluorescence complementation assay

The CDS of DRT111 and SF1 were cloned by Gateway technology in the pUGW2 and pUGW0 vectors (Nakagawa et al., 2007) to guarantee the downstream fusion of the C-terminal YFP region and upstream fusion of N-terminal YFP region, respectively. Primers are listed in Supplemental Table S2. *Nicotiana tabacum* leaf protoplasts were prepared and transfected according to Pedrazzini et al. (1997). 40 µg of DNA for each construct was introduced in 1×10^6^ protoplasts using PEG-mediated transfection. Following 16h incubation in the dark at 25°C, the cells were imaged with an Inverted Z.1 microscope (Zeiss, Germany) equipped with a Zeiss LSM 700 spectral confocal laser-scanning unit (Zeiss, Germany). Samples were excited with a 488 nm, 10 mW solid laser with emission split at 505 nm for YFP and excited with a 555 nm, 10 mW solid laser with emission split at 551 nm for chlorophyll detection

### RNA sequencing analysis

For RNA deep sequencing, total RNA was extracted from dry seeds and DNAse treated using RNAeasy plant kit (Qiagen, Hilden, Germany). Three biological replicates per genotype (Columbia-0 and *drt111-2*) were used. Library construction was performed using the Illumina TruSeq RNA Sample Preparation Kit (Illumina, SanDiego, CA, USA) prior to sequencing in single (2×100, ∼45.000.000 total reads/sample) on Illumina platform Hiseq 2500. The sequencing service was provided by Genomix4life (http://www.genomix4life.com) at Laboratory of Molecular Medicine and Genomics (University of Salerno, Italy). Raw sequences are deposited in NCBI Sequence Read Archive, bioproject PRJNA557116. Prior to further analysis, a quality check was performed on the raw sequencing data by using FastQC (https://www.bioinformatics.babraham.ac.uk/projects/fastqc/), then low quality portions of the reads were removed with BBDuk (sourceforge.net/projects/bbmap/). The minimum length of the reads after trimming was set to 35 bp and the minimum base quality score to 25. The high quality reads were aligned against the *Arabidopsis thaliana* reference genome sequence (Araport11) with STAR aligner (version 2.5.0c, Doblin et al., 2013). FeatureCounts (version 1.4.6-p5, Liao et al., 2013) was used together with the most recent *Arabidopsis thaliana* annotation to calculate gene expression values as raw read counts. Normalized TMM and FPKM values were calculated. All the statistical analyses were performed with *R* with the packages HTSFilter (Rau et al., 2013) and edgeR (Robinson et al., 2010). The first step was the removal of not expressed genes and the ones showing too much variability. The HTSFilter package was chosen for this scope, which implements a filtering procedure for replicated transcriptome sequencing data based on a Jaccard similarity index. The “Trimmed Means of M-values”(TMM) normalization strategy was used. The filter was applied to the different experimental conditions in order to identify and remove genes that appear to generate an uninformative signal. The overall quality of the experiment was evaluated, on the basis of the similarity between replicates, by a Principal Component Analysis (PCA) using the normalized gene expression values as input. The differential expression analysis was performed to identify the genes that are differentially expressed in all comparisons. Only genes with ǀlog2(fold-change)ǀ> 0.21 and FDR equal or lower than 0.05 were considered as Differentially Expressed Genes (DEGs).

In order to identify the number of different splicing events the software rMATS (V 3.2.5, Shen et al., 2014) was used. Prior to further analysis, the high quality reads were aligned against the Arabidopsis thaliana genome using Araport11 as reference with STAR aligner (version 2.5.0c), with Local Mapping option due to the restrictions in the splicing software. An FDR filter of <=0.05 was used to detect significant differences in splicing events between Col-0 and drt111. The bioinformatics analysis was performed by Sequentia Biotech (http://www.sequentiabiotech.com). For the DEGs and significantly different splicing events, a Gene Ontology Enrichment Analysis (GOEA) was performed to identify the most enriched Gene Ontology (GO) categories across the down- and up-regulated genes (P value < 0.05 and FDR <0.05) following the method described in Tian et al. 2017.

### Accession Numbers

The genes used in this study are: DRT111/SFPS (At1g30480), SUA (At3g54230), PIF1/PIL5 (At2g20180), ABI3 (At3g24650), SF1 (At5g51300), SOM (At1g03790), phyB (At2g18790).

## Acknowledgments

We are thankful to Prof. Wim Soppe, Dr. Lucio Conti and Dr. Eirini Kaiserli for providing seeds. We acknowledge Mr. Rosario Nocerino and Gaetano Guarino for excellent technical assistance. This research was carried out with funding from the Italian Ministry of University and Research, GenHORT (PON02_00395_3215002), GenoPOM-PRO (PON02_00395_3082360) and from Ministry of Agricultural, Food and Forestry Policies, BIOTECH projects. Paola Punzo was the recipient of a fellowship by Regione Campania, POR Campania FSE 2014/2020 Program “Sviluppo di processi innovativi e di prodotti di qualità per il benessere dei consumatori e la valorizzazione del comparto agroalimentare campano”.

## Supplemental data

**Supplemental Materials and Methods**

**Supplemental Figure S1.** DRT111 protein alignment.

**Supplemental Figure S2.** Sub-cellular localization of DRT111 protein.

**Supplemental Figure S3.** Identification of *DRT111* knockout mutants and over-expressing plants.

**Supplemental Figure S4.** Analysis of enriched Gene Ontology categories and validation of RNA-seq data.

**Supplemental Figure S5.** Germination of phyB-9, *pifq, pil5-1, pil5-3* and *phyB* in the presence of ABA.

**Supplemental Table S1.** List of DEGs in *drt111-2* seeds previously identified as *ABI3* targets.

**Supplemental Table S2.** List of primers used in this study.

**Supplemental Data Set1.** Genes differentially expressed in *drt111-2* seeds and Gene Ontology Enrichment Analysis.

**Supplemental Data Set2.** Alternative splicing events defective in *drt111-2* seeds.

**Supplemental Data Set3.** Gene Ontology Enrichment Analysis of alternative splicing events defective in *drt111-2* seeds.

